# Association between vitamin D deficiency and exercise capacity in patients with CKD, a cross-sectional analysis

**DOI:** 10.1101/2020.10.26.350546

**Authors:** Emma L Watson, Thomas J Wilkinson, Tom F O’Sullivan, Luke A Baker, Douglas W Gould, Soteris Xenophontos, Matthew PM Graham-Brown, Rupert W Major, Carl Jenkinson, Martin Hewison, Andrew Philp, Alice C Smith

## Abstract

Evidence is growing for a role of vitamin D in regulating skeletal muscle mass, strength and functional capacity. Given the role the kidneys play in activating total vitamin D, and the high prevalence of vitamin D deficiency in Chronic Kidney Disease (CKD), it is possible that deficiency contributes to the low levels of physical function and muscle mass in these patients. This is a secondary cross-sectional analysis of previously published interventional study, with *ex vivo* follow up work. 34 CKD patients at stages G3b-5 (eGFR 25.5 ± 8.3ml/min/1.73m2; age 61 ± 12 years) were recruited, with a sub-group (n=20) also donating a muscle biopsy. Vitamin D and associated metabolites were analysed in plasma by liquid chromatography tandem-mass spectroscopy and correlated to a range of physiological tests of muscle size, function, exercise capacity and body composition. The effects of 1α,25(OH)_2_D3 supplementation on myogenesis and myotube size was investigated in primary skeletal muscle cells from vitamin D deficient donors. *In vivo*, there was no association between total or active vitamin D and muscle size or strength, but a significant correlation with 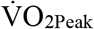 was seen with the total form. *Ex vivo*, 1α,25(OH)_2_D3 supplementation reduced IL-6 mRNA expression, but had no effect upon proliferation, differentiation or myotube diameter. This early preliminary work suggests that vitamin D deficiency is not a prominent factor driving the loss of muscle mass in CKD, but may play a role in reduced exercise capacity.

## Introduction

Patients with chronic kidney disease (CKD) commonly experience skeletal muscle wasting, reduced exercise capacity and lower levels of physical function (1–3). These appear early in the disease process (4) and are associated with adverse clinical outcomes and reduced quality of life (5–10). The factors driving loss of muscle mass and physical function are not yet fully understood but are likely to be multifactorial with large heterogeneity. Gaps in our understanding have meant that there are currently no viable therapies to protect or restore muscle mass and physical function in CKD.

The classical effects of vitamin D focus around calcium homeostasis and bone health, but it is becoming increasingly accepted that it may also have a role in skeletal muscle function (11, 12) and exercise capacity (13). Studies involving both humans and animals have shown that vitamin D deficiency is associated with muscle atrophy affecting predominately type II fibres (14, 15), which can be reversed following vitamin D supplementation (16). Vitamin D deficiency is also associated with an increased number of falls (17), which in some cases can be prevented with supplementation (18, 19). Community-based cross-sectional studies have shown vitamin D deficiency is associated with reduced measures of physical functioning such as gait speed and rising from a chair (20, 21). However, studies in both healthy and clinical populations have not always demonstrated improvements in physical functioning following vitamin D supplementation (22–24). Therefore, the role of vitamin D in the maintenance or improvement of physical function requires further examination.

Vitamin D obtained from sunlight or through dietary sources is relatively inactive and must be converted to the active form, of which the final step occurs in the kidney by 1α-hydroxylase. Given the impairment of kidney function, vitamin D deficiency is highly prevalent in CKD patients (25). Despite this, there is limited and conflicting data regarding the association between vitamin D and physical function in the CKD population. One study of CKD patients failed to find an association between the active metabolite, 1,25-dihydroxyvitamin D3 (1α,25(OH)_2_D3), and muscle function (26), whilst others in both non-dialysis CKD (27) and end-stage renal disease (28) have reported associations between 1α,25(OH)_2_D3 muscle strength and physical functioning.

Mature skeletal muscle cells are terminally differentiated, and by themselves, are capable of limited repair and regeneration. Therefore for repair to occur, cells are reliant on a population of stem cells, termed satellite cells, that support repair and regeneration through a process called myogenesis (29). Myogenesis is thought to be dysfunctional in CKD (29) and may contribute to skeletal muscle wasting. A role is also emerging for vitamin D in skeletal muscle repair (30) where it has been shown to influence satellite cell proliferation and differentiation (31, 32). Therefore, it is possible that vitamin D deficiency contributes to atrophy through inhibition of myogenesis in CKD, but this is yet to be investigated.

Vitamin D status is generally based upon the analysis of the inactive total form of vitamin D (25OHD). However, related metabolites have also been shown to be clinically important (33), demonstrating the importance to also consider the vitamin D metabolome alongside total vitamin D, which by itself provides only a limited view of vitamin D status.

The aims of this study were: 1) to perform *in vivo* analysis to determine the relationship between serum vitamin D and its metabolites and skeletal muscle mass and function in patients with CKD not requiring dialysis; and 2) to determine *ex vivo* the effect of 1α,25(OH)_2_D3 supplementation on myoblast proliferation, differentiation, and hypertrophy using human-derived skeletal muscle cells isolated from CKD vitamin D deficient donors. We hypothesised that plasma 1α,25(OH)_2_D3 would be associated with muscle size, strength and exercise capacity and that supplementation of vitamin D deficient cells with 1α,25(OH)_2_D3 would increase myotube size.

## Material and Methods

### Patients and study design

This study was a cross-sectional observational design. Patients in this report are from two separate cohorts. Physical function data is taken from the ExTra CKD study (34) (ISRCTN: 36489137), whilst patients who donated biopsies used in the *ex vivo* study took part in the Explore CKD study (ISRCTN: 18221837). Sample size was based upon available data from these participants. All patients were recruited from nephrology outpatient clinics at Leicester General Hospital, UK between December 2013 – April 2017. Exclusion criteria were age <18years, pregnancy, disability that prevented patients from undertaking exercise, insufficient command of English, or an inability to give informed consent. Ethical approval was given by the National Research Ethics Committee (13/EM/0344; 15/EM/0467). All patients gave written informed consent and the trial was conducted in accordance with the Declaration of Helsinki.

### Physiological assessments

#### Muscle size

Muscle size was determined using two methods: (i) quadriceps volume of the right leg measured by Magnetic Resonance Imaging (MRI) (34) acquired using a 3T Siemens Skyra HD MRI scanner in the axial plane using a T1 turbo spin-echo sequence and (ii) rectus femoris cross-sectional area of the right leg measured by 2-D B-mode ultrasound. These techniques have previously been described by our group (35).

#### Muscle strength

Quadriceps strength was assessed by leg extension exercise using a 5-Repetition Maximum (5-RM) test (34). Prediction equations were then used to estimate 1-Repetition Maximum (1-RM) (36).

#### Exercise capacity

Patients underwent the incremental shuttle walk test (ISWT) (34) during which patients walked along a 10m course in time with externally paced beeps that become progressively quicker until volitional fatigue. This is a valid and reliable method to determine peak exercise capacity (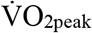) (37). Patients also underwent an incremental Cardiopulmonary Exercise Test to measure 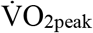 performed on an electrically-braked cycle ergometer (34).

#### Physical Function

Physical function was determined using the sit-to-stand 60 (STS60) test, a surrogate marker of muscular endurance (37).

#### Body composition

Body fat percentage and appendicular lean mass (ALM), was estimated using multi-frequency bioelectrical impedance analysis (BIA) (InBody 370, CA,USA). This device has been validated against dual-energy x-ray absorptiometry (38).

### Blood sampling and Vitamin D metabolite analysis

Venous blood samples were taken from 38 CKD patients (Table 1) into a plain tube and left undisturbed at room temperature for 30min to allow the blood to clot. The blood was then centrifuged at 1500 g for 10min at 4°C. Resulting serum was collecting and stored at −80°C until subsequent analysis. Serum concentrations of vitamin D metabolites were analysed by liquid chromatography-tandem mass spectrometry (LC-MS/MS) as previously described (39). Briefly, 200μl serum was extracted prior to analysis by protein precipitation followed by supportive liquid-liquid extraction. Analysis was performed on a Waters Acquity UPLC coupled to a Waters Xevo TQ-XS mass spectrometer. Analysis was carried out in multiple reaction monitoring (MRM) for the following analytes: 25-hydroxyvitamin D3 (25OHD3), 3-epi-25-hydroxyvitamin D3 (3-epi-25OHD3), 24,25-dihydroxyvitamin D3 (24,25(OH)_2_D3), 1,25-dihydroxyvitamin D3 (1α,25(OH)_2_D3) and 25-hydroxyvitamin D2 (25OHD2). The LC-MS/MS method for vitamin D quantification was validated for serum analysis as previously described for accuracy, precision, recovery and matrix effects (39). Vitamin D metabolites were purchased from Supleco Sigma Aldrich. LC-MS grade methanol and water were purchased from Greyhound Chromatography and Thermo Fisher respectively. Supportive liquid-liquid extraction plates were purchased from Phenomenex.

**Table 1.**
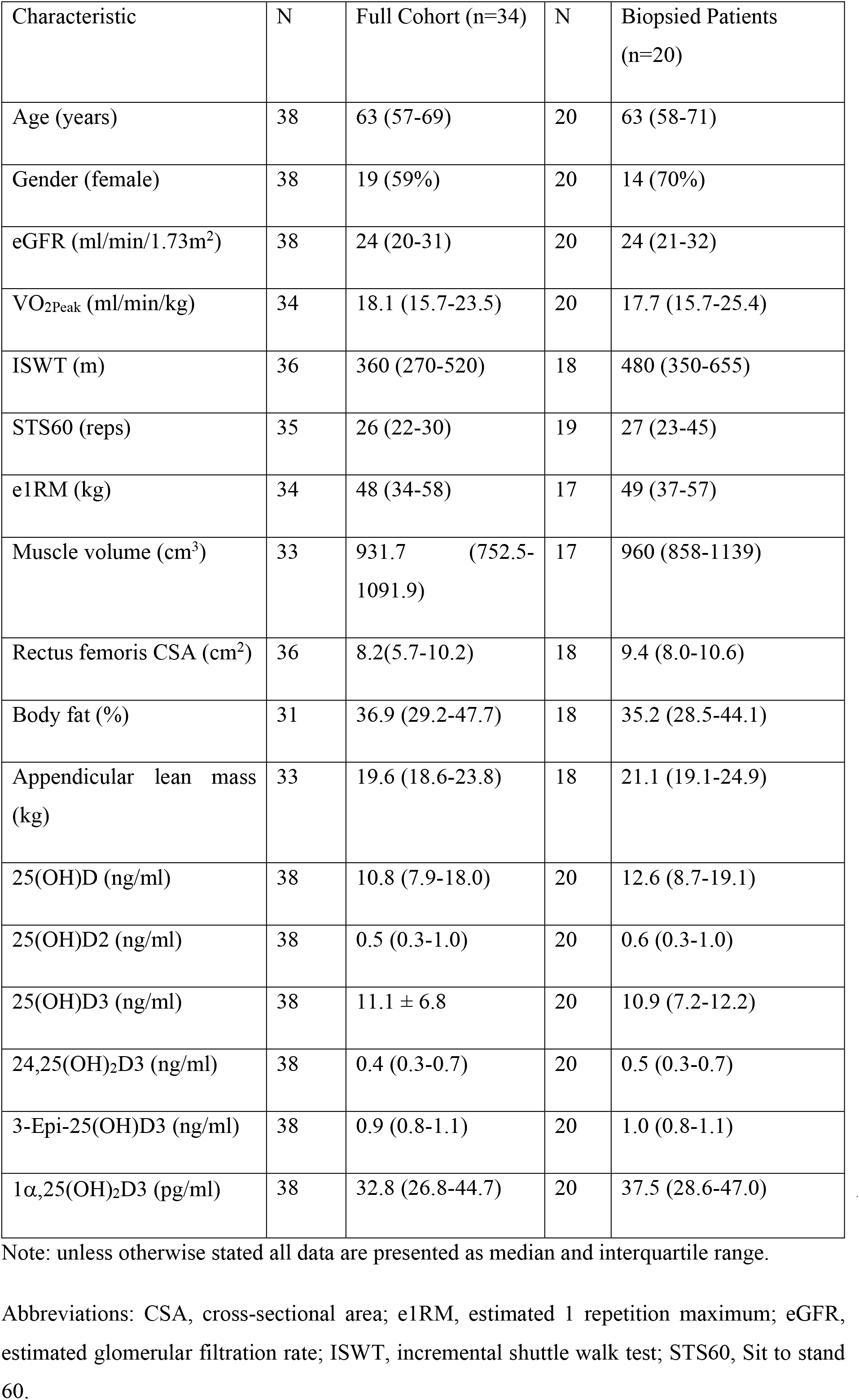
Patient characteristics for the *in vivo* study

| Characteristic | N | Full Cohort (n=34) | N | Biopsied Patients (n=20) |
| --- | --- | --- | --- | --- |
| Age (years) | 38 | 63 (57-69) | 20 | 63 (58-71) |
| Gender (female) | 38 | 19 (59%) | 20 | 14 (70%) |
| eGFR (ml/min/1.73m<sup>2</sup>) | 38 | 24 (20-31) | 20 | 24 (21-32) |
| VO<sub>2Peak</sub> (ml/min/kg) | 34 | 18.1 (15.7-23.5) | 20 | 17.7 (15.7-25.4) |
| ISWT (m) | 36 | 360 (270-520) | 18 | 480 (350-655) |
| STS60 (reps) | 35 | 26 (22-30) | 19 | 27 (23-45) |
| e1RM (kg) | 34 | 48 (34-58) | 17 | 49 (37-57) |
| Muscle volume (cm<sup>3</sup>) | 33 | 931.7 (752.5-1091.9) | 17 | 960 (858-1139) |
| Rectus femoris CSA (cm<sup>2</sup>) | 36 | 8.2(5.7-10.2) | 18 | 9.4 (8.0-10.6) |
| Body fat (%) | 31 | 36.9 (29.2-47.7) | 18 | 35.2 (28.5-44.1) |
| Appendicular lean mass (kg) | 33 | 19.6 (18.6-23.8) | 18 | 21.1 (19.1-24.9) |
| 25(OH)D (ng/ml) | 38 | 10.8 (7.9-18.0) | 20 | 12.6 (8.7-19.1) |
| 25(OH)D2 (ng/ml) | 38 | 0.5 (0.3-1.0) | 20 | 0.6 (0.3-1.0) |
| 25(OH)D3 (ng/ml) | 38 | 11.1 ± 6.8 | 20 | 10.9 (7.2-12.2) |
| 24,25(OH)<sub>2</sub>D3 (ng/ml) | 38 | 0.4 (0.3-0.7) | 20 | 0.5 (0.3-0.7) |
| 3-Epi-25(OH)D3 (ng/ml) | 38 | 0.9 (0.8-1.1) | 20 | 1.0 (0.8-1.1) |
| 1α,25(OH)<sub>2</sub>D<sub>3</sub> (pg/ml) | 38 | 32.8 (26.8-44.7) | 20 | 37.5 (28.6-47.0) |

Note: unless otherwise stated all data are presented as median and interquartile range.
Abbreviations: CSA, cross-sectional area; e1RM, estimated 1 repetition maximum; eGFR, estimated glomerular filtration rate; ISWT, incremental shuttle walk test; STS60, Sit to stand 60.

### Muscle biopsy collection

Vastus lateralis muscle biopsies were taken from 20 patients using the micro biopsy technique after an overnight fast (40). Biopsy specimens from five of these patients deemed to be vitamin D deficient (25(OH)D <20ng/ml) were also used to establish primary cultures as described below in which the effect of vitamin D repletion could be more closely studied. After dissection of any visible fat and connective tissue, samples were placed into liquid nitrogen (RNA extraction) or 5mL ice-cold Hams F10 media containing 1% penicillin streptomycin and 1% Gentamycin (cell culture).

### Satellite cell isolation procedure and cell treatments

Muscle tissue was washed in HamsF10 (containing 1% penicillin streptomycin and 1% Gentamycin), minced into small fragments and enzymatically digested in two incubations with collagenase IV (1mg/mL), Bovine Serum Albumin (BSA) (5mg/mL) and trypsin (500μl/mL) at 37°C with gentle agitation. The resultant supernatant was added to Foetal Bovine Serum (FBS), strained through a 70μm nylon filter and centrifuged at 800 g for 7min. The cells were washed in Hams F10 with 1% penicillin streptomycin and 1% Gentamycin and pre-plated on uncoated 9cm^2^ petris in 3mL growth media (GM; Hams F10 Glutamax, 20% FBS, 1% Penicillin Streptomycin, 1% fungazone) for 3h. The cell suspension was then moved to collagen I coated 25cm^2^ flasks and kept at 37°C under humidified 95% air and 5% CO_2_ until cells had achieved approximately 70% confluence. For experiments, cells were plated at a density of 3×10^4^ and grown until 70% confluent. For experiments using myoblasts, cells were exposed to either high dose of exogenous 1α,25(OH)_2_D_3_ (100nmol), low dose (10nmol), or control vehicle (95% EtOH), and proliferation rates determined after 72h. For experiments using myotubes, GM was replaced with differentiation medium (DM; DMEM 4.5g/L glucose, 1% Penicillin Streptomycin, 10% horse serum) for five days by which time multinucleated muscle fibres had formed. Cells were again exposed to either high dose of exogenous 1α,25(OH)_2_D3 (100nmol), low dose (10nmol), or control vehicle (95% EtOH) to investigate effects differentiation, determined after five days (immunofluorescence and PCR).

### Proliferation assay

Carboxyfluorescein succinimidyl ester (CFSE) dye (Thermo Fisher, UK) was added to human skeletal muscle cells in suspension (1ml HBSS) at a concentration of 5μm and incubated at 37°C for 20min. Staining was quenched by the addition of five volumes of GM and incubated at 37°C for a further 5min. Cells were rinsed and seeded onto collagen I coated 6-well plates and collected 72h later. Cells were then re-suspended in Phosphate Buffered Saline (PBS) and analysed using a FACSCELESTA instrument (BD Biosciences). Data were analysed using FlowJo 10.2 (FlowJo LLC, USA).

### Immunofluorescence

Cells were fixed in 4% paraformaldehyde for 20min at room temperature, washed three times with PBS and blocked and permeabilized in PBS containing 5% goat serum and 0.25% Triton X-100 for 1h. Cells were incubated with rabbit anti-desmin primary antibody (1/400; cell signalling) at 4°C overnight, washed three times in PBS, and incubated with Alexa Flour 488-labelled goat anti-rabbit IgG (1/400; Thermo Fisher) for 2h at room temperature. DAPI (100ng/mL) was used to visualise the nuclei. Ten random fields were acquired per condition using a FLoid imaging system (Thermo Fisher) and images analysed using ImageJ. Myotube diameter was assessed at three points on each cell. Fusion indexes were defined by the number of DAPI positive nuclei within myotubes (desmin positive cell containing 3 or more nuclei) divided by the total number of DAPI positive nuclei.

### Quantitative RT-PCR

Total RNA was extracted from skeletal muscle tissue (10mg wet weight) and primary cells using Trizol^®^ (Invitrogen, UK) and 1μg RNA was reverse transcribed to cDNA using an AMV reverse transcription system (Promega, Madison, WI, USA). Primers, probes and internal controls for all genes were supplied as Taqman gene expression assays (Applied Biosystems, Warrington, UK) Vitamin D receptor: Hs01045843_m1, Myogenin: Hs01072232_m1, MyoD: Hs02330075_g1, Myf5:Hs00929416_g1, Pax7:Hs00242962_m1, MAFbx: Hs00369714_m1, MuRF-1: Hs00822397_m1, IL-6: Hs00985639_m1, MCP-1: Hs00234140_m1, TNF-α: Hs01113624_g1, Myostatin: Hs00976237_m1, MYHC1: Hs00428600_m1, MYHC2: Hs00430042_m1, MyHC3: Hs01074230_m1, MYHC7: Hs01110632_m1, MYHC8: Hs00267293_m1 and 18s:Hs99999901_s1 was used as an internal control. All reactions were carried out in a 20 μl volume, 1 μl cDNA, 10 μl 2X Taqman Mastermix, 8 μl water, 1 μl primer/probe on an Agilent Biosystem Light Cycler with the following conditions, 95°C 15s, followed by 40X at 95°C for 15s and 60°C for 1 min. The Ct values from the target gene were normalized to 18s and expression levels calculated according to 2^-ΔΔ Ct^ method to determine fold changes.

### Statistical analysis

All data were tested for normality using the Shapiro-Wilk test. If data were not normally distributed, analysis was performed on log-transformed data, or non-parametric tests were used as appropriate. Partial correlations, controlled for age and gender, were performed to determine the relationship between vitamin D metabolites and physiological outcomes. Spearman’s or Pearson’s bivariate correlations were performed as appropriate to determine the relationship between vitamin D metabolites and skeletal muscle gene expression data. The LC/MS-MS lower detection limit for 1α,25(OH)_2_D3 analysis is 32pg/ml. A sensitivity analysis was performed removing cases that fell below this threshold. For all PCR data, one-way repeated measures ANOVA was performed on the ΔCT values with predetermined pairwise comparisons (vehicle vs 10nm 1α,25(OH)_2_D3, vehicle vs 100nm 1α,25(OH)_2_D3 and 10 vs 100nm 1α,25(OH)_2_D3. Differences in outcome measures for patients classified as ‘deficient’ vs ‘insufficient’ were analysed by linear regression with age and gender added into the model as covariates. For tissue culture work data were expressed as fold change compared to the vehicle control condition (2^-ΔΔCT^). Effect sizes were estimated using Cohens d or eta squared (n^2^) statistic as appropriate (d; interpreted small ≥ 0.20, medium ≥ 0.50, large ≥ 0.80; n^2^; interpreted small ≥ 0.01, medium ≥ 0.06, large ≥ 0.14). Missing data was analysed using Little’s test, to test the assumption of missing completely at random (MCAR). This showed that missing data was MCAR and so a complete case analysis was performed All statistical analyses were performed using IBM SPSS 25 software (IBM, Chicago, IL). Statistical significance was accepted as P<0.05.

## Results

### Patient characteristics and vitamin D status

Patient characteristics for the *in vivo* study can be found in Table 1. In summary, median age was 63 (57-69 years), 19/34 patients were females, median eGFR was 24 (20-31 ml/min/1.73m^2^). Of these, 28/34 (82%) patients were vitamin D deficient (25(OH)D: <20ng/ml), and a further 6 (18%) were insufficient (25(OH)D: 21-29ng/ml). No patients exhibited sufficient vitamin D levels (25(OH)D: >30ng/ml) according to the Endocrine Society guideline criteria (41). Using the National Kidney Foundation Kidney Disease Outcome Quality Initiative (NKF KDOQI) guidelines, 4/34 (12%) patients were severely deficient (25(OH)D: <5ng/ml), 20/34 (59%) had a mild deficiency (25(OH)D: 5-15ng/ml), and 10/34 (29%) were insufficient (25(OH)D: 16-30ng/ml). No patients were classified as having sufficient vitamin D levels (25(OH)D: >30ng/ml) (42). Regardless of which criteria are used, all patients fell below the cut-offs defined for intervention (41).

### *In vivo* investigation of the association between vitamin D and its metabolites and measures of muscle mass and physical function

#### Associations with physiological assessments

Correlations between vitamin D metabolites and physiological assessments can be found in Table 3. Positive, although small, correlations were seen between total vitamin D and both VO_2Peak_ (rho = 0.41, p = 0.04) and STS60 performance (rho = 0.45, p = 0.02), as well as between total vitamin D and ISWT (rho = 0.37 p = 0.06), e1-RM (rho = 0.36, p = 0.07), and percentage body fat (rho = −0.39, p = 0.05). We saw a moderate association between the active form of vitamin D and 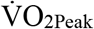 (rho = 0.53, p = 0.005), e-1RM (rho = 0.50, p = 0.008) and STS 60 performance (rho = 0.49, p = 0.01). No meaningful association was seen with RF-CSA (rho = 0.35, p = 0.08). However, all these relationships disappeared when individuals with values <32pg/ml were removed from the analysis (n = 19). When the cohort was split for ‘deficiency’ compared to ‘insufficiency’ based upon NKF-KDOQI guidelines, there was a significant difference in 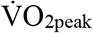 between the groups (deficiency: 17.4 (15.0-20.5) vs insufficiency: 22.3 (17.7-28.4ml/min/kg; p = 0.006) but not for performance in the ISWT (deficiency: 350 (262-495) vs insufficiency: 395 (345-672m; p = 0.16). Characteristics for patients within these groups can be found in supplementary table 1.

#### Associations with skeletal muscle gene expression

Characteristics of the 20 patients who donated muscle biopsies used in this *ex vivo* analysis can be found in Table 2. Partial correlations can be found in Table 4. A negative association was seen between 25OHD and Activin type II receptor (rho −0.69, p = 0.03), MuRF-1 (rho −0.75, p = 0.01) and MAFbx (rho −0.79, p = 0.006). No other correlations were observed.

**Table 2.** Patient characteristics for *ex vivo* biopsy analysis and cell culture experiments.

| Characteristic | Donors used in biopsy PCR analysis | Donors used in primary cell culture experiments |
| --- | --- | --- |
| n | 20 | 5 |
| Age (years) | 58 (57-63) | 57 (52-58) |
| Gender (female) | 4 (80%) | 4 (80%) |
| eGFR (ml/min/1.73m<sup>2</sup>) | 23 (19-29) | 23 (19-29) |
| 25(OH)D (ng/ml) | 10.0 (9.2-12.0) | 10.0 (9.2-12.0) |
| VO<sub>2Peak</sub> (ml/min/kg) | 16.7 (15.9-24.5) | 16.1 (15.7-27.0) |
| ISWT (m) | 540 (370-560) | 575 (495-635) |
| STS60 (reps) | 27 (26-39) | 28 (27-38) |
| e1RM (kg) | 48 (44-53) | 53 (50-54) |
| Muscle volume (cm<sup>3</sup>) | 973 (955-1039) | 960 (916-1026) |
| Rectus femoris CSA (cm<sup>2</sup>) | 8.7 (7.5-9.5) | 9.0 (8.2-9.6) |
| Body fat (%) | 34 (30-43) | 39.3 (35.9-44.2) |

Note: unless otherwise stated all data are presented as median and interquartile range.
Abbreviations: CSA, cross-sectional area; e1RM, estimated 1 repetition maximum; eGFR, estimated glomerular filtration rate; ISWT, incremental shuttle walk test; STS60, Sit to stand 60.

**Table 3.** Bivariate correlations between serum vitamin D and vitamin D metabolites and physiological outcome measures Data are correlation coefficients with p values in brackets. * denotes p<0.05

| Physiological outcome measure | 25(OH)D | 25OHD2 | 25OHD3 | 24,25(OH)<sub>2</sub>D3 | 3-Epi-25OHD3 | 1α,25(OH)<sub>2</sub>D3 | 1α,25(OH)<sub>2</sub>D3 (>32pg/ml<sup>§</sup>) |
| --- | --- | --- | --- | --- | --- | --- | --- |
| eGFR (ml/min/1.72m<sup>2</sup>) | 0.017 (p = 0.99) | -0.15 (p = 0.55) | -0.008 (p = 0.97) | 0.22 (p = 0.28) | 0.47 (p = 0.01) | 0.14 (p = 0.48) | -0.13 (p = 0.66) |
| VO<sub>2Peak</sub> (ml/min/kg) | **0.41\* (p = 0.04)** | 0.38 (p = 0.14) | 0.36 (p = 0.70) | **0.40\* (p = 0.04)** | 0.25 (p = 0.21) | **0.53\* (p = 0.005)** | 0.25 (p = 0.39) |
| ISWT (m) | 0.37 (p = 0.06) | 0.08 (p = 0.75) | 0.36 (p = 0.07) | **0.40\* (p = 0.04)** | **0.41\* (p = 0.04)** | 0.25* (p = 0.20) | -0.04 (p = 0.89) |
| e-1RM (kg) | 0.36 (p = 0.07) | -0.26 (p = 0.30) | 0.36 (p = 0.07) | 0.33 (p = 0.10) | **0.43\* (p = 0.03)** | **0.50\* (p = 0.008)** | 0.31 (p = 0.28) |
| RF-CSA (cm<sup>2</sup>) | 0.28 (p = 0.16) | -0.45 (p = 0.07) | 0.28 (p = 0.16) | 0.33 (p = 0.11) | **0.47\* (p = 0.02)** | 0.35 (p = 0.08) | 0.04 (p = 0.89) |
| Quadriceps volume (cm<sup>3</sup>) | 0.03 (p = 0.89) | **-0.50\* (p = 0.04)** | 0.04 (p = 0.83) | 0.16 (p = 0.43) | 0.22 (p = 0.27) | 0.24 (p = 0.24) | -0.04 (p = 0.89) |
| Appendicular lean mass (kg) | 0.27 (p = 0.17) | -0.60 (p = 0.10) | 0.30 (p = 0.13) | 0.31 (p = 0.12) | **0.47\* (p = 0.02)** | 0.36 (p = 0.10) | 0.12 (p = 0.67) |
| Body fat (%) | -0.39 (p = 0.05) | -0.15 (p = 0.58) | -0.38 (p = 0.06) | -0.22 (p = 0.28) | -0.21 (p = 0.29) | -0.07 (p = 0.70) | -0.25 (p = 0.39) |
| STS60 (reps) | **0.45\* (p = 0.02)** | 0.20 (p = 0.45) | **0.43\* (p = 0.03)** | 0.36 (p = 0.07) | 0.15 (p = 0.45) | **0.49\* (p = 0.01)** | 0.47 (p = 0.09) |

Abbreviations: e-1RM, estimated 1-repetition maximum; eGFR, estimated glomerular filtration rate; ISWT, incremental shuttle walk test; RF-CSA, rectus femoris cross-sectional area. STS60, sit to stand 60.
<sup>§</sup> Cases <32pg/ml removed as below the LC/MS-MS lower detection limit

**Table 4.** Partial correlations between serum vitamin D and vitamin D metabolites and expression levels of genes involved in processes of maintenance of muscle mass.

| Gene | 25(OH)D | 25OHD2 | 25OHD3 | 24,25(OH)<sub>2</sub>D3 | 3-Epi-25OHD3 | 1α,25(OH)<sub>2</sub>D3 |
| --- | --- | --- | --- | --- | --- | --- |
| Myostatin | 0.06 (p = 0.83) | -0.48 (p = 0.16) | 0.17 (p = 0.49) | -0.04 (p = 0.87) | -0.05 (p = 0.86) | 0.06 (p = 0.84) |
| Activin type II receptor | -0.08 (p = 0.79) | **-0.69\* (p = 0.03)** | 0.13 (p = 0.57) | -0.04 (p = 0.89) | -0.03 (p = 0.92) | 0.19 (p = 0.53) |
| MuRF-1 | 0.07 (p = 0.82) | **-0.75\* (p = 0.01)** | 0.12 (p = 0.64) | 0.12 (p = 0.69) | -0.03 (p = 0.90) | 0.29 (p = 0.32) |
| MAFbx | -0.06 (p = 0.83) | **-0.79\* (p = 0.006)** | -0.02 (p = 0.92) | 0.02 (p = 0.92) | 0.06 (p = 0.84) | 0.24 (p = 0.41) |
| MyoD | -0.30 (p = 0.31) | -0.49 (p = 0.14) | -0.17 (p = 0.45) | -0.12 (p = 0.68) | -0.02 (p = 0.93) | -0.21 (p = 0.47) |
| Myf5 | 0.12 (p = 0.69) | -0.23 (p = 0.51) | 0.10 (p = 0.68) | 0.18 (p = 0.55) | 0.11 (p = 0.70) | -0.11 (p = 0.70) |
| Myogenin | -0.16 (p = 0.60) | -0.59 (p = 0.07) | -0.04 (p = 0.87) | -0.10 (p = 0.72) | 0.08 (p = 0.97) | -0.04 (p = 0.89) |
| Pax7 | -0.13 (p = 0.67) | -0.63 (p = 0.05) | -0.11 (p = 0.65) | 0.02 (p = 0.92) | -0.05 (p = 0.85) | 0.17 (p = 0.56) |
| TNF-α | -0.33 (p = 0.26) | 0.002 (p = 0.99) | -0.28 (p = 0.24) | -0.27 (p = 0.35) | -0.38 (p = 0.20) | -0.11 (p = 0.71) |
| IL-6 | -0.36 (p = 0.23) | -0.55 (p = 0.10) | -0.28 (p = 0.24) | -0.35 (p = 0.24) | -0.33 (p = 0.26) | 0.02 (p = 0.93) |
| MCP-1 | -0.18 (p = 0.54) | -0.55 (p = 0.09) | 0.02 (p = 0.93) | -0.10 (p = 0.73) | -0.01 (p = 0.95) | -0.29 (p = 0.33) |

Data are correlation coefficients with p values in brackets. \* denotes p<0.05
Abbreviations: IL-6, Interleukin-6; MAFbx, muscle atrophy F box; MCP-1, monocyte chemoattractant protein-1; MuRF-1. Muscle RING finger 1; myf5, myogenic factor 5; myoD, myoblast determination protein; TNF-α , tumor necrosis factor alpha.

#### *Ex vivo* investigation of the effect of Vitamin D on human derived skeletal muscle cells

Characteristics of those patients (n=5) used in this investigation can be found in Table 2.

#### Effects on inflammation, protein degradation and myogenesis

Doses of both 10 and 100nm 1α,25(OH)_2_D3 reduced IL-6 mRNA expression in myotubes 2-fold compared to the vehicle condition (p = 0.03, d = 1.5; p = 0.02, d = 0.9 respectively), which were both large effects. However, there was no significant effect of either dose on TNF-α (p = 0.35; d = 0.18; Figure 1). There was a trend for 1α,25(OH)_2_D3 to reduce expression of myostatin by 1.6-fold (10nm) and 2-fold (100nm) compared to the vehicle, but this was only a small effect (p = 0.07, η^2^ = 0.47). There was also a trend to reduce expression of MuRF-1 by 1.8-fold (10nm) and 1.1-fold (100nm; p = 0.08, η^2^ = 0.57). No effect was seen of either dose on MAFbx expression (p = 0.32, η^2^ = 0.25; Figure 2). When the expression of the myogenic regulatory factors was determined, there was a trend for 1α,25(OH)_2_D3 to reduce myogenin expression by 1.4-fold (10nm) and 2-fold (100nm) (p = 0.08, η^2^ = 0.46), but MyoD expression was unchanged (p = 0.10, η^2^ = 0.44). There was no effect of 1α,25(OH)_2_D3 on pax7 expression (p = 0.42, η^2^ = 0.21; Figure 3). 10nm 1α,25(OH)_2_D3 was seen to significantly reduce expression of MYHC1 compared to the vehicle condition by 5-fold (p = 0.04, d = 0.57) and a similar 5-fold reduction was seen for 100nm (p = 0.09, d = 1.0). An effect of 10nm 1α,25(OH)_2_D3 was also seen on MyHC8 expression, which was reduced by 2.5-fold (p = 0.03, d = 0.53), a similar reduction was seen with 100nm (p = 0.07, d = 1.01; Figure 4). No effect of either dose was seen on MYHC2 (p = 0.12, η^2^ = 0.41), MYHC3 (p = 0.07, η^2^ = 0.58) or MYHC7 (p = 0.23, η^2^ = 0.31) mRNA expression. There was no effect of 1α,25(OH)_2_D3 at either dose on mRNA expression of the Vitamin D receptor (p = 0.39, η^2^ = 0.13).

**Figure 1.**
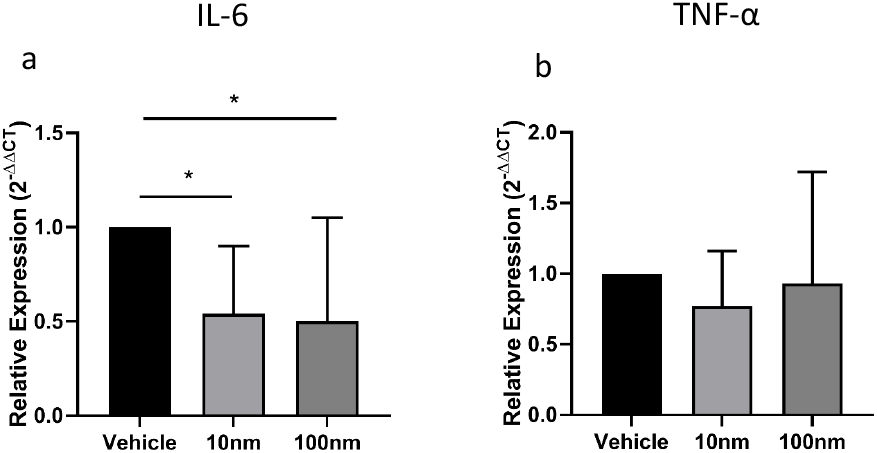
Effect of 10nm and 100nm 1α,25(OH)_2_D3 vs vehicle on inflammatory cytokine mRNA expression. a) IL-6; b) TNF-α. Real time PCR data presented as 2^-ΔΔCT^ relative to vehicle. * denotes P<0.05 vs baseline.

**Figure 2.**
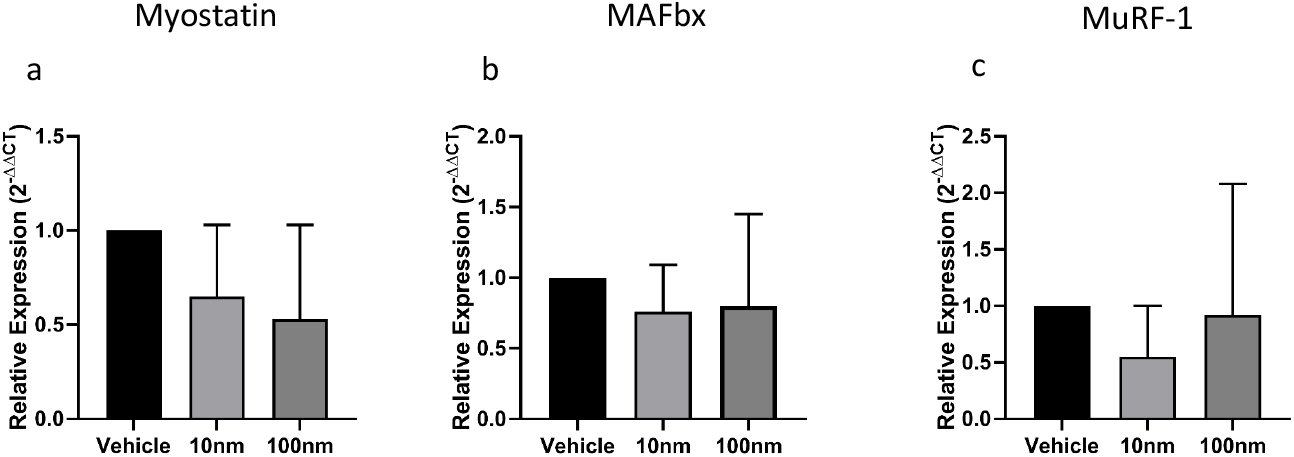
Effect of 10nm and 100nm 1α,25(OH)_2_D3 vs vehicle on mRNA expression of proteins relating to muscle protein breakdown. a) Myostatin; b) MAFbx; c) MuRF-1.

**Figure 3.**
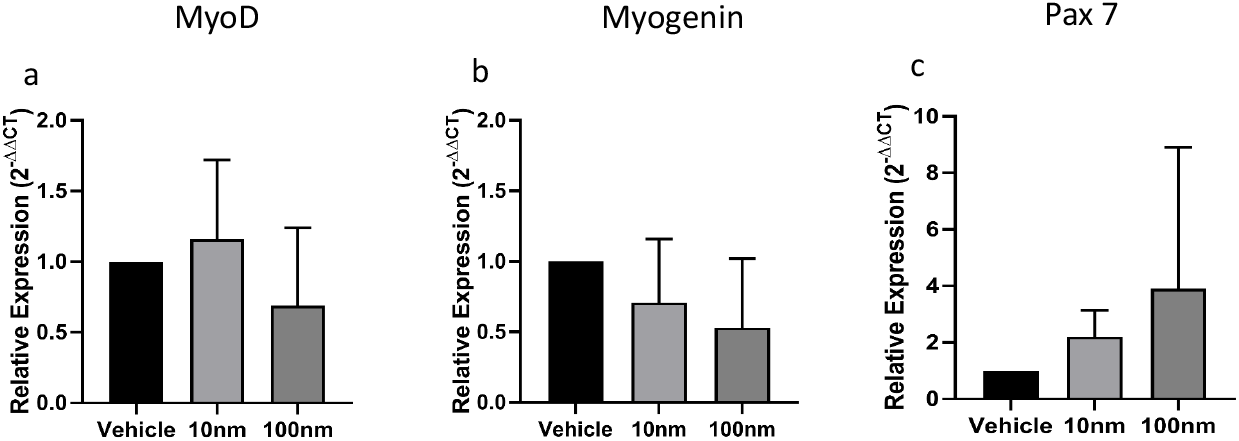
Effect of 10nm and 100nm 1α,25(OH)_2_D3 vs vehicle on mRNA expression of proteins relating to myogenesis. a) MyoD; b) myogenin; c) Pax7.

**Figure 4.**
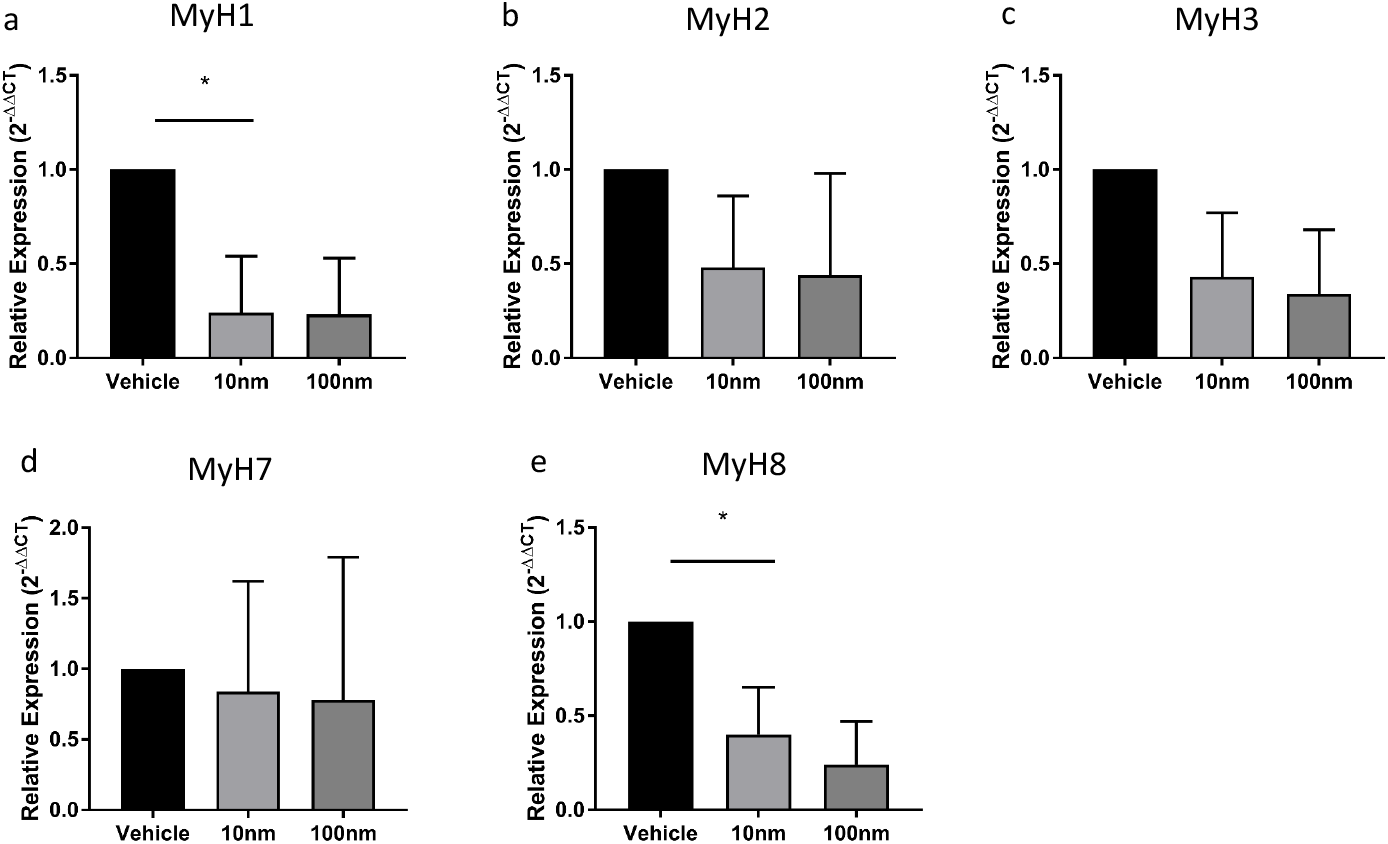
Effect of 10nm and 100nm 1α,25(OH)_2_D3 vs vehicle on mRNA expression of myosin heavy chain isoforms. a) MyH1; b) MyH2; c) MyH3; d) MyH7; e) MyH8. * denotes P<0.05 vs vehicle.

#### Effects on morphology

There was no effect of either 10nm or 100nm 1α,25(OH)_2_D3 compared to vehicle on myotube diameter (10nm = 18.2 ± 3.1 vs 100nm = 19.7 ± 4.4 vs vehicle = 16.5 ± 2.4μm; p = 0.37, η^2^ = 0.22), or fusion index (10nm = 23.0 ± 9.5 vs 100nm = 23.8 ± 12.8 vs vehicle = 22.9 ± 6.4%; p = 0.85, η^2^ = 0.05). 100nm 1α,25(OH)_2_D3 resulted in significantly fewer myotubes per field of view compared to the vehicle (p = 0.03, d = 0.84), with no differences between vehicle vs 10nm (p = 0.11, d = 0.28) or 10nm vs 100nm (p = 0.24, d = 0.38) (10nm = 4 ± 4 vs 100nm = 3 ± 2 vs vehicle = 5 ± 3 myotubes per field of view; Figure 5).

**Figure 5.**
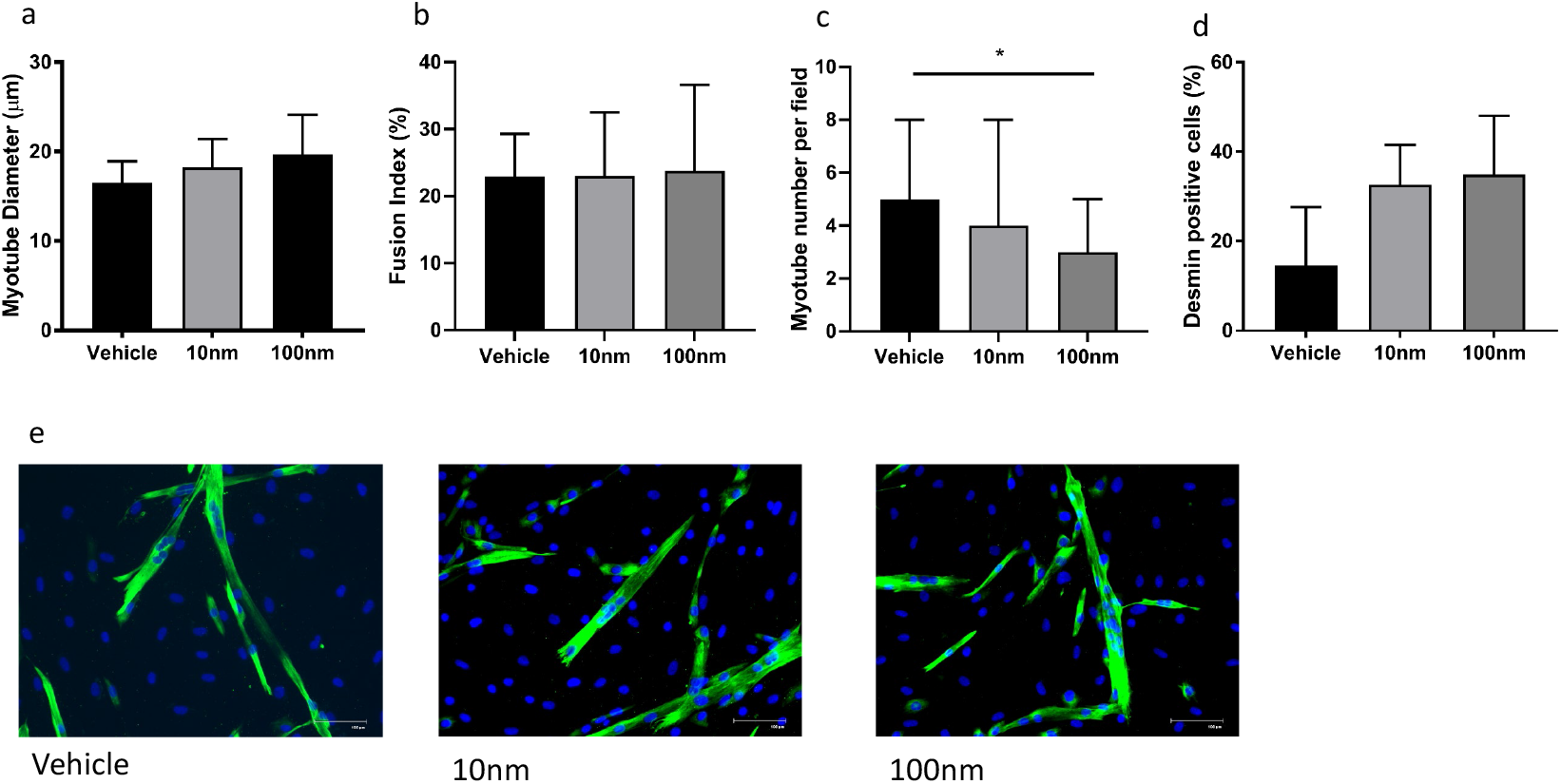
Effect of 10nm and 100nm 1α,25(OH)_2_D3 vs vehicle on myotube morphology. a) myotube diameter (μm); b) myotube number per field; c) fusion index (%); d) desmin positive cells (%); e) representative immunofluroscent images. Cells are counterstained for desmin (green) and DAPI (blue). Scale bar represents 100μm. * denotes P<0.05 vs vehicle.

#### Effect on cell proliferation

There was no effect of 1α,25(OH)_2_D3 on myoblast proliferation, with no difference in geometric mean fluorescent intensity of CFSE in response to either 10nm or 100nm dose, compared to vehicle (P = 0.77, η^2^ = 0.09).

## Discussion

The aims of this study were two-fold. Firstly, to perform *in vivo* analysis to determine the relationship between serum vitamin D and its metabolites with skeletal muscle mass and function in CKD patients not requiring dialysis and to establish if vitamin D deficiency might contribute to reduced physical function. Secondly, to determine if 1α,25(OH)_2_D3 supplementation in human-derived skeletal muscle cells established from CKD vitamin D deficient donors, could improve myoblast proliferation, differentiation, and myotube size.

Our data demonstrates a high prevalence of vitamin D deficiency, with all patients meeting the guidelines for initiation of vitamin D supplementation (25(OH)D levels: <30ng/ml). This is in line with previous reports (41, 43) and highlights the need to understand the physiological effects of such a deficiency. Data from healthy population cohorts demonstrate a relationship between vitamin D and muscle function. In particular, vitamin D levels are associated with muscular strength (44) and physical function (21), with vitamin D supplementation reducing the risk of falls in elderly populations (45). However, the influence of vitamin D status on muscle mass or size is less clear. A study in vitamin D receptor knock out mice found smaller muscle fibres compared to their sham littermates (46), but a study of nearly 700 healthy men and women was unable to find an association between total or active vitamin D and muscle mass (47). The authors concluded that the link between falls and vitamin D may be due to effects upon neuromuscular function rather than muscle mass.

There is a scarcity of data available on the role of vitamin D in muscle mass/function in the CKD population. Gordon and colleagues reported that active vitamin D was associated with gait speed, sit to stand tests, 6-minute walk test, and isokinetic muscle strength in CKD patients at stage G3-4 (27). They also saw an association with muscle size when calcium and physical activity levels were controlled for. A similar association has also been observed between total vitamin D (although not active vitamin D), muscle strength, and falls also in CKD patients not requiring dialysis (28). Further, vitamin D supplementation in CKD patients at stages G3-4 has been shown to improve measures of physical functioning (48). We observed a significant association between both the total and active forms of vitamin D with and VO_2Peak_, and performance in the STS-60. This tentatively suggests a role for vitamin D in determining exercise capacity and physical function in these patients. An association between serum vitamin D and cardiorespiratory fitness has been reported recently in the healthy population (49), which may be driven through effects upon oxidative metabolism (50, 51). Interestingly, we have recently shown in the same cohort of patients as those reported on here, reduced mitochondrial number compared to a healthy control cohort (52). It would, therefore, be important to know the effect of vitamin D on skeletal muscle mitochondrial function in these patients. It is also possible that this association of vitamin D with exercise capacity and physical function is acting through an effect upon patient’s symptom and fatigue perception rather than a direct effect upon skeletal muscle *per se*. This is an interesting observation that warrants closer investigation.

We also report some associations between vitamin D metabolites and the physiological outcome measures. However, there is limited previously published data on these relationships (53) and more research is required to understand the importance of these associations. These relatively ambiguous results highlight the need for more definitive studies to better understand the relationship between the vitamin D metabolome and physical function and muscle function and mass and the relative importance of supplementation in this group.

As it was thought vitamin D may play a role in the maintenance of muscle mass (46), we hypothesised that the addition of 1α,25(OH)_2_D3 to primary skeletal muscle cells from vitamin D deficient donors would reduce myoblast proliferation and increase myoblast differentiation and myotube size. Studies of C2C12 cells and primary skeletal muscle cells have shown that Vitamin D has anti-proliferative effects (32, 54). However, we saw no effect of either dose of 1α,25(OH)_2_D3 on myoblast proliferation rates after 72h of treatment, which has been reported previously in C2C12 cells (55), although a lower dose of 1nm was used in this study. There was also no effect of supplementation on differentiation, fusion index, desmin positivity, or myotube diameter and no change in the mRNA expression of MyoD or myogenin, which are usually required for myotube fusion. Rather, we observed a reduction in the mRNA expression of MyHC1 (type IIX) and MyHC8 (perinatal) with 10nm 1α,25(OH)_2_D3, which is in contrast to previous reports (54). We did, however, observe a decrease in the number of myotubes formed following 100nm 1α,25(OH)_2_D3, which has been reported before in both primary human skeletal muscle cells and C2C12 cells (30, 31). Therefore, the reduction in MyHC expression may just be an artefact of the reduced number of myotubes in culture. Vitamin D has also been reported to have hypertrophic effects *ex vivo* (30, 56), with 1α,25(OH)_2_D3 supplementation resulting in increases in myotube diameter. However, no evidence of 1α,25(OH)_2_D3 stimulated hypertrophy was seen here. The reasons for these discrepancies are currently unclear. A lack of an effect on hypertrophy may be the result of the absence of a change in myostatin mRNA expression. Both *in vitro* and *in vivo* studies have demonstrated that vitamin D supplementation reduces skeletal muscle myostatin mRNA expression (32, 54). It is possible that CKD has induced a higher myostatin expression in these cells (57), which is not modifiable by 1α,25(OH)_2_D3 exposure and that this higher expression level prevents hypertrophy driven by other mechanisms. The lack of any large effect on our human-derived skeletal muscle cells is in agreement with our *in vivo* data, which failed to find any associations between vitamin D and measures of muscle size or strength. This lends support to the notion that vitamin D does not directly affect skeletal muscle in these patients.

Interestingly, IL-6 mRNA expression was reduced with both doses of 1α,25(OH)_2_D3, which is in contrast to *in vivo* results, where there was no relationship seen between any form of vitamin D and skeletal muscle IL-6 mRNA expression. There has been little reported regarding the anti-inflammatory properties of vitamin D in skeletal muscle, but there is evidence from other systems (58). The effect on IL-6 was seen in the absence of any reduction of TNF-α expression, and was, therefore, unlikely to have resulted in significant anti-inflammatory affects. Given the role IL-6 is known to play in skeletal muscle wasting (59), this does warrant further investigation.

There are a few limitations of this study that should be taken into account. Firstly, this is a secondary analysis of an earlier study and was therefore not powered to detect relationships between vitamin D deficiency and physical function or muscle mass. Given the prevalence of vitamin D deficiency in these patients and the role this plays in physical function in other groups, a suitably powered study is warranted to better understand its implications. We suspect that many of the discrepancies in the *ex vivo* results presented here might be explained by differences in the models used in the experiments, human vs rodent and immortalised vs primary cell lines. The effect of vitamin D administration in human-derived skeletal muscle cells from CKD donors has not been investigated before. Given the complicated and diverse effects of CKD upon skeletal muscle physiology, it is likely that vitamin D supplementation is not sufficient to overcome more potent effects imposed by the illness. These results are only based on five patients that exhibit a large degree of variation which is likely masking real effects. A larger sample size might provide more definitive results and as such these results should only be considered preliminary. Finally, this study has performed individual metabolite analysis only. It may be interesting to apply modelling strategies that take into account the interplay between the different metabolites (60) to get a full overview of the effect of the vitamin D metabolome on physical function and muscle mass in these patients.

In conclusion, we have seen no strong evidence for a role of total or active vitamin D in determining the level of muscle size or strength in these patients. At the cellular level, our preliminary data suggests there is no effect of 1α,25(OH)_2_D3 supplementation on myoblast proliferation, differentiation or hypertrophy. We did, however, see an association between total vitamin D and VO_2Peak_ and STS60 performance which was also seen with active vitamin D before the sensitivity analysis. This suggests that vitamin D deficiency is not a prominent factor driving the loss of muscle mass in CKD, but may have a role to play in the poor exercise tolerance and low exercise capacity seen in these patients. In light of the high prevalence of vitamin D deficiency in this patient population, further investigation is warranted to understand the role of this hormone on skeletal muscle physiology in CKD.

## Acknowledgements

The authors thank all research assistants involved in data and sample collection. The research was supported by the National Institute for Health Research (NIHR) Leicester Biomedical Research Centre. The views expressed are those of the authors and not necessarily those of the NHS, the NIHR or the Department of Health. The staff and running costs of this study were part-funded by the Stoneygate Trust and by an early career grant awarded by the Society for Endocrinology to Dr Emma Watson. Dr Emma Watson was supported by Kidney Research UK (PDF2/2015). Dr Major was funded by Kidney Research UK (TF2/2015).

## Author contributions

EW, TWJ, DWG, MH, CJ, AP and AS were involved in study conception and plan. EW, TJW, SX, MGB, RM, and DWG were involved in patient recruitment and performed assessments. EW, LB, DWG, TOS, CJ, MH and AP were involved in laboratory analysis, data analysis interpretation. EW, LB, TJW, AP and AS were responsible for preparing the manuscript for submission. All authors approve this submission.

## Abbreviations

1-RM: 1 repetition maximum
1α,25(OH)_2_D3: 1,25-dihydroxyvitamin D3
24,25(OH)_2_D3: 24,25-dihydroxyvitamin D3
25OHD2: 25-hydroxyvitamin D2
25OHD3: 25-hydroxyvitamin D3
3-epi-25OHD3: 3-epi-25-hydroxyvitamin D3,
5-RM: 5 repetition maximum
ALM: Appendicular lean mass
BIA: Bioelectrical impedance analysis
BSA: Bovine serum albumin
CFSE: Carboxyfluorescein succinimidyl ester
CKD: Chronic kidney disease
DM: Differentiation medium
FBS: fetal bovine serum
eGFR: estimated glomerular filtration rate
GM: Growth medium
IL-6: Interleukin-6
ISWT: Incremental shuttle walk test
LC-MS/MS: liquid chromatography-tandem mass spectrometry
MCP-1: monocyte chemoattractant protein-1
MRI: Magnetic resonance imaging
MRM: Multiple reaction monitoring
MAFbx: Muscle atrophy F-box
MuRF-1: Muscle ring finger-1
MyHC: Myosin heavy chain
MyoD: Myoblast determination protein 1
NKF-KDOQI: National Kidney Foundation Kidney Disease Outcome Quality Initiative
PBS: Phosphate buffered saline
STS60: sit-to-stand 60
TNF-α: Tumour necrosis factor alpha
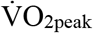: Peak oxygen uptake

## References

1. Mak RH, Ikizler AT, Kovesdy CP, Raj DS, Stenvinkel P, Kalantar-Zadeh K. Wasting in chronic kidney disease. J Cachexia Sarcopenia Muscle. 2011;2(1):9–25.

2. Segura-Orti E, Gordon PL, Doyle JW, Johansen KL. Correlates of Physical Functioning and Performance Across the Spectrum of Kidney Function. Clin Nurs Res. 2018;27(5):579–96.

3. Koufaki P, Mercer T. Assessment and monitoring of physical function for people with CKD. Adv Chronic Kidney Dis. 2009;16(6):410–9.

4. John SG, Sigrist MK, Taal MW, McIntyre CW. Natural history of skeletal muscle mass changes in chronic kidney disease stage 4 and 5 patients: an observational study. PLoS One. 2013;8(5):e65372.

5. Roshanravan B, Robinson-Cohen C, Patel KV, Ayers E, Littman AJ, de Boer IH, et al. Association between physical performance and all-cause mortality in CKD. J Am Soc Nephrol. 2013;24(5):822–30.

6. Roshanravan B, Gamboa J, Wilund K. Exercise and CKD: Skeletal Muscle Dysfunction and Practical Application of Exercise to Prevent and Treat Physical Impairments in CKD. Am J Kidney Dis. 2017;69(6):837–52.

7. Hiraki K, Yasuda T, Hotta C, Izawa KP, Morio Y, Watanabe S, et al. Decreased physical function in pre-dialysis patients with chronic kidney disease. Clin Exp Nephrol. 2013;17(2):225–31.

8. Painter P. Physical functioning in end-stage renal disease patients: update 2005. Hemodial Int. 2005;9(3):218–35.

9. Carrero JJ, Chmielewski M, Axelsson J, Snaedal S, Heimburger O, Barany P, et al. Muscle atrophy, inflammation and clinical outcome in incident and prevalent dialysis patients. Clin Nutr. 2008;27(4):557–64.

10. MacKinnon HJ, Wilkinson TJ, Clarke AL, Gould DW, O’Sullivan TF, Xenophontos S, et al. The association of physical function and physical activity with all-cause mortality and adverse clinical outcomes in nondialysis chronic kidney disease: a systematic review. Ther Adv Chronic Dis. 2018;9(11):209–26.

11. Girgis CM, Clifton-Bligh RJ, Hamrick MW, Holick MF, Gunton JE. The roles of vitamin D in skeletal muscle: form, function, and metabolism. Endocr Rev. 2013;34(1):33–83.

12. Ceglia L, Harris SS. Vitamin D and its role in skeletal muscle. Calcif Tissue Int. 2013;92(2):151–62.

13. Pandey A, Kitzman DW, Houston DK, Chen H, Shea MK. Vitamin D Status and Exercise Capacity in Older Patients with Heart Failure with Preserved Ejection Fraction. Am J Med. 2018;131(12):1515.e11-.e19.

14. Tieland M, Brouwer-Brolsma EM, Nienaber-Rousseau C, van Loon LJ, De Groot LC. Low vitamin D status is associated with reduced muscle mass and impaired physical performance in frail elderly people. Eur J Clin Nutr. 2013;67(10):1050–5.

15. Endo I, Inoue D, Mitsui T, Umaki Y, Akaike M, Yoshizawa T, et al. Deletion of vitamin D receptor gene in mice results in abnormal skeletal muscle development with deregulated expression of myoregulatory transcription factors. Endocrinology. 2003;144(12):5138–44.

16. Bhat M, Kalam R, Qadri SS, Madabushi S, Ismail A. Vitamin D deficiency-induced muscle wasting occurs through the ubiquitin proteasome pathway and is partially corrected by calcium in male rats. Endocrinology. 2013;154(11):4018–29.

17. Snijders T, Verdijk LB, Beelen M, McKay BR, Parise G, Kadi F, et al. A single bout of exercise activates skeletal muscle satellite cells during subsequent overnight recovery. Exp Physiol. 2012;97(6):762–73.

18. Pfeifer M, Begerow B, Minne HW, Suppan K, Fahrleitner-Pammer A, Dobnig H. Effects of a long-term vitamin D and calcium supplementation on falls and parameters of muscle function in community-dwelling older individuals. Osteoporos Int. 2009;20(2):315–22.

19. Bischoff HA, Stahelin HB, Dick W, Akos R, Knecht M, Salis C, et al. Effects of vitamin D and calcium supplementation on falls: a randomized controlled trial. J Bone Miner Res. 2003;18(2):343–51.

20. Bischoff-Ferrari HA, Dietrich T, Orav EJ, Hu FB, Zhang Y, Karlson EW, et al. Higher 25-hydroxyvitamin D concentrations are associated with better lower-extremity function in both active and inactive persons aged > or =60 y. Am J Clin Nutr. 2004;80(3):752–8.

21. Wicherts IS, van Schoor NM, Boeke AJ, Visser M, Deeg DJ, Smit J, et al. Vitamin D status predicts physical performance and its decline in older persons. J Clin Endocrinol Metab. 2007;92(6):2058–65.

22. Witham MD, Crighton LJ, Gillespie ND, Struthers AD, McMurdo ME. The effects of vitamin D supplementation on physical function and quality of life in older patients with heart failure: a randomized controlled trial. Circ Heart Fail. 2010;3(2):195–201.

23. Levis S, Gomez-Marin O. Vitamin D and Physical Function in Sedentary Older Men. J Am Geriatr Soc. 2017;65(2):323–31.

24. Bischoff-Ferrari HA, Dawson-Hughes B, Orav EJ, Staehelin HB, Meyer OW, Theiler R, et al. Monthly High-Dose Vitamin D Treatment for the Prevention of Functional Decline: A Randomized Clinical Trial. JAMA Intern Med. 2016;176(2):175–83.

25. Obi Y, Hamano T, Isaka Y. Prevalence and prognostic implications of vitamin D deficiency in chronic kidney disease. Dis Markers. 2015;2015:868961.

26. Heaf JG, Molsted S, Harrison AP, Eiken P, Prescott L, Eidemak I. Vitamin D, surface electromyography and physical function in uraemic patients. Nephron Clin Pract. 2010;115(4):c244–50.

27. Gordon PL, Doyle JW, Johansen KL. Association of 1,25-dihydroxyvitamin D levels with physical performance and thigh muscle cross-sectional area in chronic kidney disease stage 3 and 4. J Ren Nutr. 2012;22(4):423–33.

28. Boudville N, Inderjeeth C, Elder GJ, Glendenning P. Association between 25-hydroxyvitamin D, somatic muscle weakness and falls risk in end-stage renal failure. Clin Endocrinol (Oxf). 2010;73(3):299–304.

29. Wang XH, Du J, Klein JD, Bailey JL, Mitch WE. Exercise ameliorates chronic kidney disease-induced defects in muscle protein metabolism and progenitor cell function. Kidney Int. 2009;76(7):751–9.

30. Owens DJ, Sharples AP, Polydorou I, Alwan N, Donovan T, Tang J, et al. A systems-based investigation into vitamin D and skeletal muscle repair, regeneration, and hypertrophy. Am J Physiol Endocrinol Metab. 2015;309(12):E1019–31.

31. Girgis CM, Clifton-Bligh RJ, Mokbel N, Cheng K, Gunton JE. Vitamin D signaling regulates proliferation, differentiation, and myotube size in C2C12 skeletal muscle cells. Endocrinology. 2014;155(2):347–57.

32. Garcia LA, King KK, Ferrini MG, Norris KC, Artaza JN. 1,25(OH)2vitamin D3 stimulates myogenic differentiation by inhibiting cell proliferation and modulating the expression of promyogenic growth factors and myostatin in C2C12 skeletal muscle cells. Endocrinology. 2011;152(8):2976–86.

33. Bosworth CR, Levin G, Robinson-Cohen C, Hoofnagle AN, Ruzinski J, Young B, et al. The serum 24,25-dihydroxyvitamin D concentration, a marker of vitamin D catabolism, is reduced in chronic kidney disease. Kidney Int. 2012;82(6):693–700.

34. Watson EL, Gould DW, Wilkinson TJ, Xenophontos S, Clarke AL, Vogt BP, et al. Twelve-week combined resistance and aerobic training confers greater benefits than aerobic training alone in nondialysis CKD. Am J Physiol Renal Physiol. 2018;314(6):F1188–f96.

35. Watson EL, Greening NJ, Viana JL, Aulakh J, Bodicoat DH, Barratt J, et al. Progressive Resistance Exercise Training in CKD: A Feasibility Study. Am J Kidney Dis. 2015;66(2):249–57.

36. Brzycki M. Strength testing-predicting a one-rep max from reps to fatigue. Journal of Physical Education, Recreation and Dance. 1993;64:88.

37. Wilkinson TJP, Xenophontos SM, Gould DWP, Vogt BPP, Viana JLP, Smith ACP, et al. Test-retest reliability, validation, and “minimal detectable change” scores for frequently reported tests of objective physical function in patients with non-dialysis chronic kidney disease. Physiother Theory Pract. 2019;35(6):565–76.

38. Wilkinson TJ, Richler-Potts D, Nixon DGD, Neale J, Smith AC. Anthropometry-based Equations to Estimate Body Composition: A Suitable Alternative in Renal Transplant Recipients and Patients With Nondialysis Dependent Kidney Disease? J Ren Nutr. 2019;29(1):16–23.

39. Jenkinson C, Taylor AE, Hassan-Smith ZK, Adams JS, Stewart PM, Hewison M, et al. High throughput LC-MS/MS method for the simultaneous analysis of multiple vitamin D analytes in serum. J Chromatogr B Analyt Technol Biomed Life Sci. 2016;1014:56–63.

40. Watson EL, Viana JL, Wimbury D, Martin N, Greening NJ, Barratt J, et al. The Effect of Resistance Exercise on Inflammatory and Myogenic Markers in Patients with Chronic Kidney Disease. Front Physiol. 2017;8:541.

41. Holick MF, Binkley NC, Bischoff-Ferrari HA, Gordon CM, Hanley DA, Heaney RP, et al. Evaluation, treatment, and prevention of vitamin D deficiency: an Endocrine Society clinical practice guideline. J Clin Endocrinol Metab. 2011;96(7):1911–30.

42. Foundation NK. K/DOQI clinical practice guidelines for bone metabolism and disease in chronic kidney disease. Am J Kidney Dis. 2003;42(4 Suppl 3):S1–201.

43. Mehrotra R, Kermah D, Budoff M, Salusky IB, Mao SS, Gao YL, et al. Hypovitaminosis D in chronic kidney disease. Clin J Am Soc Nephrol. 2008;3(4): 1144–51.

44. Grimaldi AS, Parker BA, Capizzi JA, Clarkson PM, Pescatello LS, White MC, et al. 25(OH) vitamin D is associated with greater muscle strength in healthy men and women. Med Sci Sports Exerc. 2013;45(1):157–62.

45. Rejnmark L. Effects of vitamin d on muscle function and performance: a review of evidence from randomized controlled trials. Ther Adv Chronic Dis. 2011;2(1):25–37.

46. Girgis CM, Cha KM, Houweling PJ, Rao R, Mokbel N, Lin M, et al. Vitamin D Receptor Ablation and Vitamin D Deficiency Result in Reduced Grip Strength, Altered Muscle Fibers, and Increased Myostatin in Mice. Calcif Tissue Int. 2015;97(6):602–10.

47. Marantes I, Achenbach SJ, Atkinson EJ, Khosla S, Melton LJ, 3rd, Amin S. Is vitamin D a determinant of muscle mass and strength? J Bone Miner Res. 2011;26(12):2860–71.

48. Taskapan H, Baysal O, Karahan D, Durmus B, Altay Z, Ulutas O. Vitamin D and muscle strength, functional ability and balance in peritoneal dialysis patients with vitamin D deficiency. Clin Nephrol. 2011;76(2):110–6.

49. Marawan A, Kurbanova N, Qayyum R. Association between serum vitamin D levels and cardiorespiratory fitness in the adult population of the USA. Eur J Prev Cardiol. 2019;26(7):750–5.

50. Ashcroft SP, Bass JJ, Kazi AA, Atherton PJ, Philp A. The vitamin D receptor regulates mitochondrial function in C2C12 myoblasts. Am J Physiol Cell Physiol. 2020;318(3):C536–c41.

51. Ryan ZC, Craig TA, Folmes CD, Wang X, Lanza IR, Schaible NS, et al. 1alpha,25-Dihydroxyvitamin D3 Regulates Mitochondrial Oxygen Consumption and Dynamics in Human Skeletal Muscle Cells. J Biol Chem. 2016;291(3):1514–28.

52. Watson EL, Baker LA, Wilkinson TJ, Gould DW, Graham-Brown MPM, Major RW, et al. Reductions in skeletal muscle mitochondrial mass are not restored following exercise training in patients with chronic kidney disease. Faseb j. 2020;34(1): 1755–67.

53. Hassan-Smith ZK, Jenkinson C, Smith DJ, Hernandez I, Morgan SA, Crabtree NJ, et al. 25-hydroxyvitamin D3 and 1,25-dihydroxyvitamin D3 exert distinct effects on human skeletal muscle function and gene expression. PLoS One. 2017;12(2):e0170665.

54. van der Meijden K, Bravenboer N, Dirks NF, Heijboer AC, den Heijer M, de Wit GM, et al. Effects of 1,25(OH)2 D3 and 25(OH)D3 on C2C12 Myoblast Proliferation, Differentiation, and Myotube Hypertrophy. J Cell Physiol. 2016;231(11):2517–28.

55. Stio M, Celli A, Treves C. Synergistic effect of vitamin D derivatives and retinoids on C2C12 skeletal muscle cells. IUBMB Life. 2002;53(3):175–81.

56. Girgis CM, Clifton-Bligh RJ, Turner N, Lau SL, Gunton JE. Effects of vitamin D in skeletal muscle: falls, strength, athletic performance and insulin sensitivity. Clin Endocrinol (Oxf). 2014;80(2):169–81.

57. Verzola D, Procopio V, Sofia A, Villaggio B, Tarroni A, Bonanni A, et al. Apoptosis and myostatin mRNA are upregulated in the skeletal muscle of patients with chronic kidney disease. Kidney Int. 2011;79(7):773–82.

58. Liu W, Zhang L, Xu HJ, Li Y, Hu CM, Yang JY, et al. The Anti-Inflammatory Effects of Vitamin D in Tumorigenesis. Int J Mol Sci. 2018;19(9).

59. Haddad F, Zaldivar F, Cooper DM, Adams GR. IL-6-induced skeletal muscle atrophy. J Appl Physiol (1985). 2005;98(3):911–7.

60. Beentjes CHL, Taylor-King JP, Bayani A, Davis CN, Dunster JL, Jabbari S, et al. Defining vitamin D status using multi-metabolite mathematical modelling: A pregnancy perspective. J Steroid Biochem Mol Biol. 2019;190:152–60.

